# Novel Genetic Risk factors for Asthma in African American Children: Precision Medicine and The SAGE II Study

**DOI:** 10.1101/043018

**Authors:** MJ White, O Risse-Adams, P Goddard, MG Contreras, J Adams, D Hu, C Eng, SS Oh, A Davis, K Meade, E Brigino-Buenaventura, MA Lenoir, K Bibbins-Domingo, M Pino-Yanes, E Burchard

## Abstract

**Background:** Asthma, an inflammatory disorder of the airways, is the most common chronic disease of children worldwide. There are significant racial/ethnic disparities in asthma prevalence, morbidity and mortality among U.S. children. This trend is mirrored in obesity, which may share genetic and environmental risk factors with asthma. The majority of asthma biomedical research has been performed in populations of European decent.

**Objective:** We sought to identify genetic risk factors for asthma in African American children. We also assessed the generalizability of genetic variants associated with asthma in European and Asian populations to African American children.

**Methods:** Our study population consisted of 1227 (812 asthma cases, 415 controls) African American children with genome-wide single nucleotide polymorphism (SNP) data. Logistic regression was used to identify associations between SNP genotype and asthma status.

**Results:** We identified a novel variant in the *PTCHD3* gene that is significantly associated with asthma (rs660498, p = 2.2 x10^−7^) independent of obesity status. Approximately 5% of previously reported asthma genetic associations identified in European populations replicated in African Americans.

**Conclusions:** Our identification of novel variants associated with asthma in African American children, coupled with our inability to replicate the majority of findings reported in European Americans, underscores the necessity for including diverse populations in biomedical studies of asthma.

## Introduction

Asthma is a chronic inflammatory disorder of the airways that affects at least 334 million people around the world, (Global Asthma Network 2014). In the U.S., racial and ethnic disparities in asthma are among the most significant among chronic diseases (Oh et al. 2016). Specifically, African American children are twice as likely as European Americans to suffer from asthma (Akinbami et al. 2014; Burchard 2014) and to die from complications of the disease (Gorina 2012).

Asthma is a multi-factorial disease likely affected by genetic and environmental factors. The presence of a strong genetic influence on disease susceptibility is evident, with heritability estimates from twin studies ranging from 63% to 92% (Fagnani et al. 2008; McGeachie et al. 2013; Nieminen et al. 1991). However, to date, genome-wide association studies (GWAS) of asthma have uncovered only a small number of loci with small to modest effects (Ober and Yao 2011). Together, these loci explain a small portion of asthma heritability (McGeachie et al. 2013). Despite the attention and resources allocated to genetic research in the past decade, globally diverse populations are severely understudied. To date, 96% of all GWAS have been performed in populations of European ancestry; specifically, only 1.9% of pulmonary studies have included members of racial/ethnic minority groups (Burchard et al. 2015; Bustamante et al. 2011). Notably, the few studies that have been performed in non-European populations have revealed the existence of ethnic-specific loci not identified in other populations (Torgerson et al. 2011).

The disparity in asthma prevalence is mirrored in obesity, in which African American children have higher prevalence of obesity than European Americans (Guerrero et al. 2016). Previous studies report overlapping risk factors between asthma and obesity. Although the co-occurrence of asthma and obesity is common, the exact causal relationship between the two phenotypes remains unclear (Thomsen et al. 2007). Moreover, among African Americans, the association of obesity with asthma is stronger than in other racial/ethnic groups (Joseph et al. 2016). Genetic associations with asthma may potentially be mediated by obesity status. Pleiotropic effects between obesity and asthma at the single nucleotide polymorphism (SNP) or gene-level may partly explain the association between these two conditions (Gonzalez et al. 2014; Melen et al. 2013; Melen et al. 2010; Wang et al. 2015). The majority of asthma genetic studies have not explored whether obesity mediates the association between identified genetic variants and asthma status (Akhabir and Sandford 2011; Hoffjan et al. 2003). Identifying loci related to asthma and obesity in minority populations has the potential of revealing genetic mechanisms underlying asthma severity and the response to therapy (McGarry et al. 2015; Sheehan and Phipatanakul 2015).

To address these gaps in clinical and biomedical research, we performed a genome-wide association study of asthma in African American. We then evaluated the impact of obesity as a mediator on significant associations between genetic variants and asthma. We also investigated whether genetic risk factors for asthma, previously identified in European and Asian populations, were generalizable to African Americans.

## Material and Methods

### Study Population

SAGE II (Study of African Americans, Asthma, Genes, and Environments) is the largest ongoing gene-environment interaction study of asthma in African American children in the U.S. SAGE II includes detailed clinical, social and environmental data on asthma and asthma-related conditions with corresponding environmental exposure, demographic, and psychosocial information. Full details of the SAGE II study protocols have been described in detail elsewhere (Borrell et al. 2013; Nishimura et al. 2013; Thakur et al. 2013). Briefly, SAGE II was initiated in 2006 and recruited participants with and without asthma through a combination of clinic‐ and community-based recruitment centers in the San Francisco Bay Area. Institutional review boards of participant centers approved the study and all participants or, for participants 17 or younger, their parents, provided written informed consent. Participants 17 or younger also provided age-appropriate assent. Asthma cases were defined as individuals with a history of physician-diagnosed asthma and asthma controller or rescue medication use within the last two years, and report of symptoms. Participants were eligible if they were 8-21 years of age and self-identified as African American and had four African American grandparents. Study exclusion criteria included the following: 1) any smoking within one year of the recruitment date; 2) 10 or more pack-years of smoking; 3) pregnancy in the third trimester; 4) history of lung diseases other than asthma (cases) or chronic illness (cases and controls). The SAGE II study enrolled 1556 (920 cases and 636 controls) from 2006 to August 2013.

### Genotyping and Quality Control

DNA was isolated from whole blood collected from 1381 SAGE II participants (817 cases and 564 controls) at the time of study enrollment using the Wizard^®^ Genomic DNA Purification Kits (Promega, Fitchburg, WI). Samples were genotyped with the Affymetrix Axiom^®^ LAT1 array (World Array 4, Affymetrix, Santa Clara, CA), which includes 817,810 SNPs. This array was designed to capture genetic variation in African-descent populations such as African Americans and Latinos (Hoffmann et al. 2011).

Genotype call accuracy and Axiom array-specific quality control metrics were assessed and applied according to the protocol described in further detail in Online Resource 1. Briefly, quality control inclusion criteria consisted of genotyping efficiency > 95%, Hardy-Weinberg Equilibrium (HWE) p > 10^−6^, and minor allele frequency (MAF) > 5%. Cryptic relatedness was also assessed to ensure that samples were effectively unrelated (no closer than third degree relatives). After quality control procedures, 797,128 single nucleotide polymorphisms (SNPs) were available for analysis in a total of 1,227 (812 asthma cases, 415 controls) individuals with complete measurements of global African ancestry, age, and sex.

### Calculation of BMI Categories

Body mass index (BMI) was calculated using height and weight according to the standardized formula 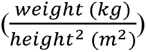 (Center for Disease Control and Prevention 2014). BMI categories were defined as follows: Obese (BMI ≥ 30) and Non-obese (BMI < 30) for participants 20 and over). For individuals under 20 years of age, BMI measurements were first converted to age‐ and gender-specific BMI percentiles (BMI-pct) before assignment to BMI categories using the following criteria: Obese (BMI-pct > 95) and Nonobese (BMI-pct < 95) (Center for Disease Control and Prevention 2014; McGarry et al. 2015). BMI information was only available for 1080 study participants (88%).

### Genetic Ancestry Estimation

The genetic ancestry of each participant was determined using the ADMIXTURE software package (Alexander et al. 2009) and modeled assuming two ancestral populations (African and European) using the HapMap Phase III data from the YRI and CEU populations as reference (International HapMap et al. 2010) (Figure 1).

**Figure 1.**
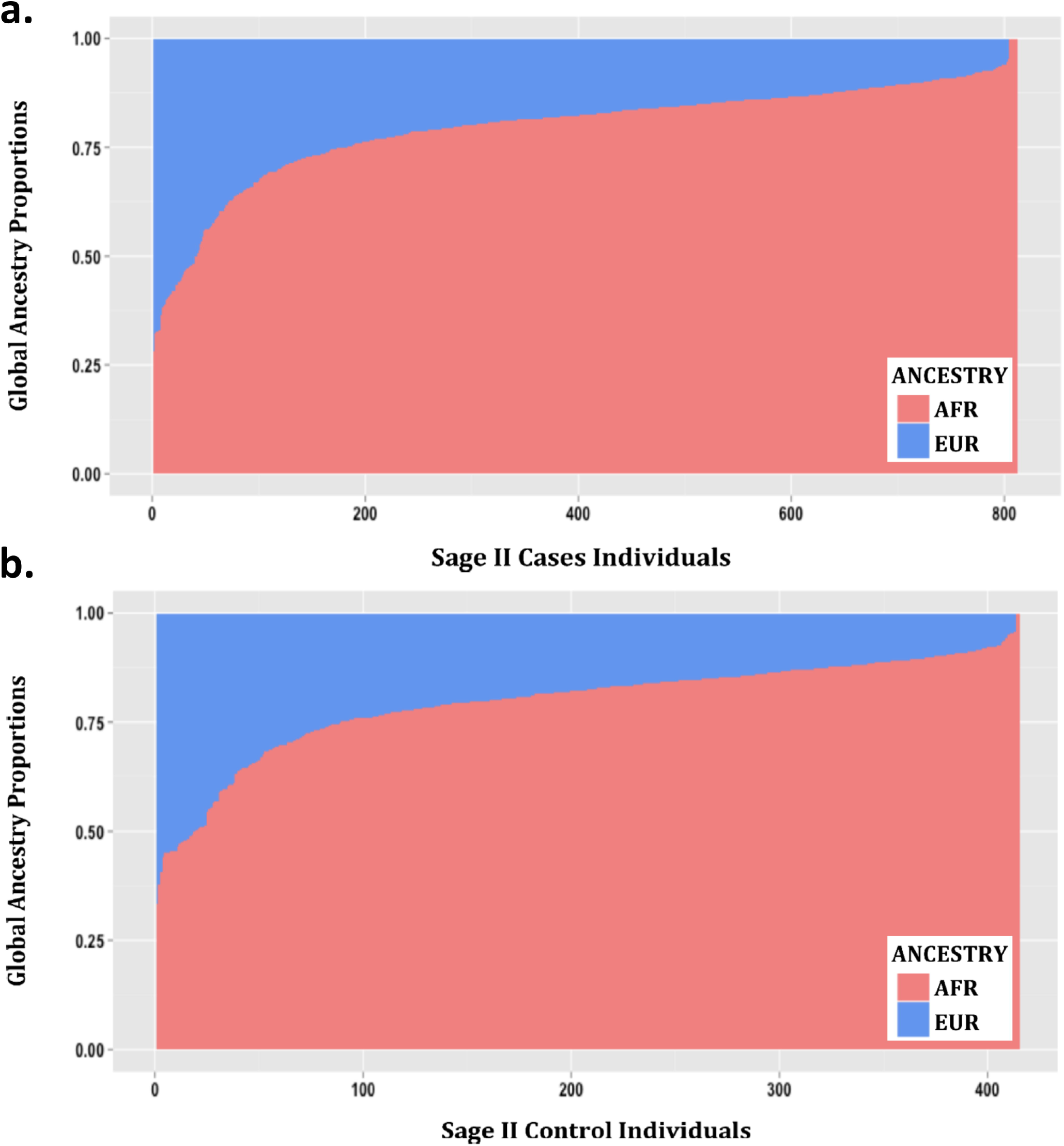
Admixture proportions assessed with ADMIXTURE for the SAGE II participants included in this study: a.) Cases, b.) Controls. Each vertical bar represents one individual and colors display the proportions of European (blue) and African (red) ancestry.

### Calculation of Population-specific Genome-wide Significance Threshold

A population-specific genome-wide significance threshold of 4.3 x 10^−7^ was derived from the number of effectively independent SNPs in our dataset, calculated according to the protocol published by Sobota *et al.* (2015). A suggestive threshold was set at p < 10^−6^ for association results.

### Statistical Analysis

#### Comparison of Demographic Variables

Demographic characteristics were compared between asthma cases and controls (Table 1). If normally distributed, continuous variables were compared using the Student’s t-test; if variables were non-normally distributed, the Wilcoxon Rank Sum test was performed. Dichotomous variables were assessed using the Chi-square test. Differences in global African ancestry proportions between cases and controls were assessed using the difference in proportion test (Acock 2014; Wang 2000). All tests were performed using STATA version 12 (StataCorp. 2011).

**Table 1.** Distribution of Demographic Characteristics for SAGE II Participants

| Characteristics |  | Cases (812) | Controls (415) | P-Value<sup>1</sup> |
| --- | --- | --- | --- | --- |
| Age, yrs<sup>2</sup> |  | 13.6 (10.9-16.8) | 16.4 (12.8-18.8) | < 0.001 |
| Gender (% Female) |  | 47.2 (43.7-50.6) | 56.1 (51.4-60.9) | 0.003<sup>#</sup> |
| Body Mass Index<sup>3</sup> | Obese | 268 | 70 | 0.002 |
|  | Non-Obese | 523 | 219 |  |
| African Ancestry<sup>2</sup> |  | 82.0 (77.0-87.0) | 82.0 (76.0-87.0) | 0.810 |

<sup>1</sup>P-values presented are from the Wilcoxon Rank Sum test with the exception of BMI (Chi-square test)
<sup>2</sup> Median (interquartile range)
<sup>3</sup>Participants > 20 years of age with BMI ≥ 30 were classified as obese; participants < 20 years of age with BMI ≥ 95<sup>th</sup> age- and gender-specific percentile were classified as obese. All other participants were classified as non-obese; frequencies in cases and controls are presented.
<sup>#</sup>P-value is calculated from difference in proportions test

#### Association Testing

Logistic regression was performed to assess the relationship between SNP genotype and asthma status. Genetic variants were coded to assess the additive effect of the minor allele (genotype coding: 0 = homozygous major, 1 = heterozygous, 2 = homozygous minor). All regression models were adjusted for age, sex, and global African ancestry. All testing was performed using the PLINK1.9 software package (Chang et al. 2015; Purcell 2015). Quantile-quantile (QQ) (Figure 2) and Manhattan (Figure 3) plots were generated using the “qqman” package (Turner 2014) in the R statistical software environment (R Development Core Team 2010). Significant and suggestive associations were further evaluated to determine if SNP effects were mediated by obesity status. Mediation analysis was performed using the “mediation” package in the R statistical software suite (Imai et al. 2010; Tingley 2014) to determine the proportion of the total association due to an indirect effect of SNP genotype on asthma status via obesity (Online Resource 2, Table S1). Non-parametric bootstrap resampling (n=1000) was used to generate 95% confidence intervals and p-values for predicted mediation estimates.

**Figure 2.**
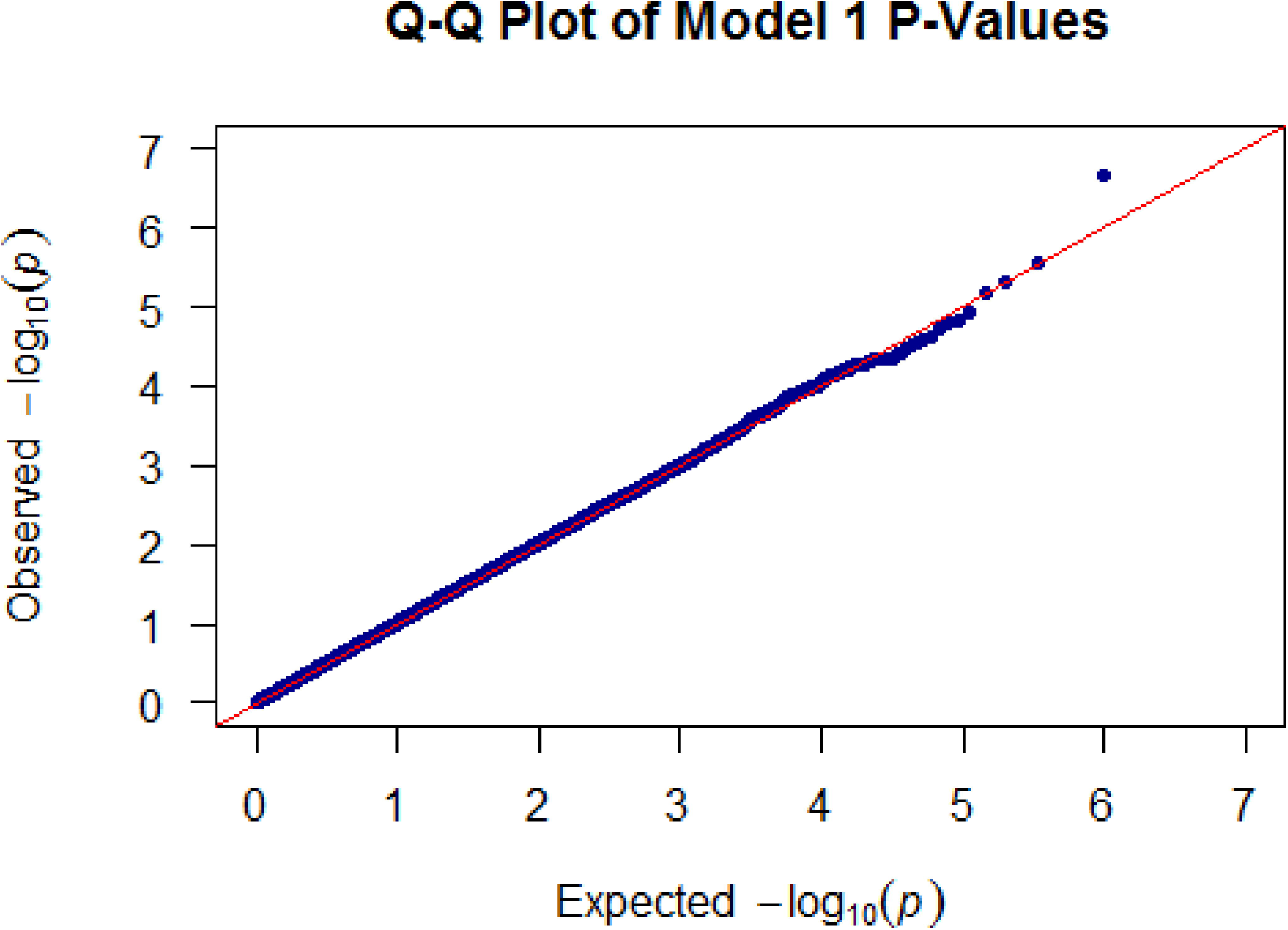
Quantile-quantile plot of the GWAS of asthma.

**Figure 3.**
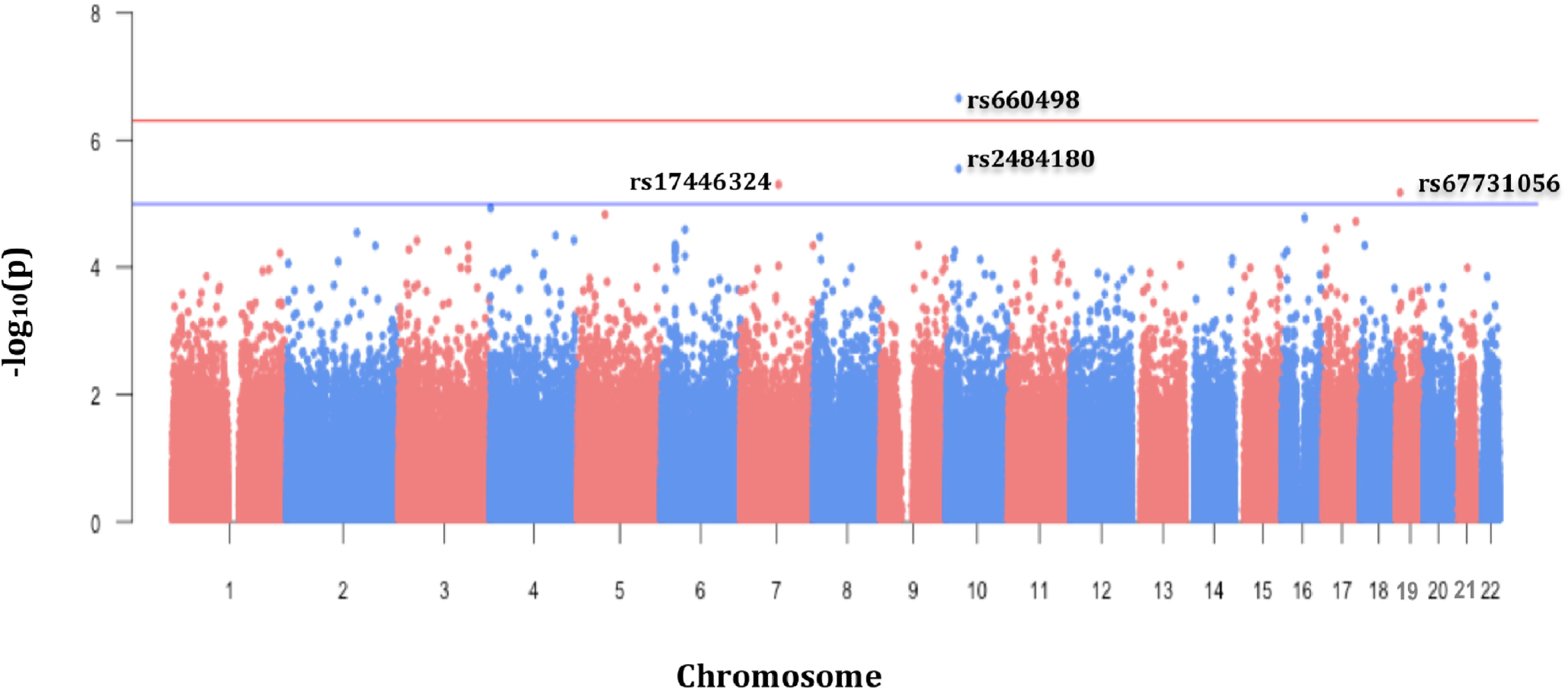
Manhattan plot of the GWAS of asthma. In the x-axis the chromosomal position of each SNP is represented and in the y-axis the–log_10_ transformation of the association p-value for each SNP.

#### Replication of Previous Asthma Associations

We identified 59 previously reported SNP associations with asthma from 17 separate studies by assessing the NHGRI-EBI Catalog of Published GWAS (Burdett 2014; Welter et al. 2014) using the search term “asthma” and a significance threshold of 5 x 10^−8^ (Online Resource 2, Table S2). It is important to note that this number includes only studies focused specifically on asthma and not distinct, but related, phenotypes such as drug response, airflow obstruction, etc.). Of the 59 variants, 39 were genotyped in our study population (Online Resource 2, Table S2). The remaining 20 variants were imputed using the IMPUTE2 (Howie et al. 2011; Howie et al. 2009) software package, and haplotypes from all available populations in the 1000 Genomes Project Phase 1 as a reference panel (Genomes Project et al. 2010), obtained with SHAPEIT (Delaneau et al. 2012). SNPs with an IMPUTE2 information score < 0.3 were excluded from analysis. A total of 59 variants remained for replication analysis following quality control procedures (Online Resource 2, Table S2 – S3).

Allelic dosage data from the imputed SNPs were analyzed for association with asthma status using logistic regression under an additive model; genotypes were coded to measure the effect of the minor allele as previously described. Regression covariates included age, sex, and global African Ancestry. The significant threshold for replication was p ≤ 0.05. Regression analyses were performed using the PLINK1.9 software package, as previously mentioned.

### Functional annotation of associated variants

For variants with association p-values that were genome-wide significant or suggestive, we performed a scan of functionality using HaploReg v4.1 (Ward and Kellis 2012) to query the empirical data from the ENCODE project and several expression quantitative trait loci (eQTL) databases. We also used the online bioinformatics data mining tool, SNPInfo, to determine if associated SNPs were predicted to affect biological functions (Xu and Taylor 2009).

## Results

### Study Population

Demographic information for the SAGE II individuals included in this study (n=1227) is presented in Table 1. The median age in control individuals (median = 16.4) was significantly higher than among asthma cases (median = 13.6) (p < 0.001). The percentage of females was also higher in controls (56.1%) relative to cases (47.2%) (p = 0.003). Obesity was significantly associated with asthma status (p = 0.002). However, no significant differences in global African ancestry were observed between asthma cases and controls (p = 0.81, Figure 1).

### Asthma GWAS

Our GWAS results did not show evidence of inflation due to population stratification (Figure 2). We identified two variants on chromosome 10 in the Patched Domain-Containing Protein 3 (*PTCHD3*) gene region, that were significantly (rs660498, OR = 1.62; p = 2.20 x 10^−7^), and suggestively (rs2484180, OR = 1.54; p = 2.84 x 10^−6^) associated with increased asthma susceptibility (Table 2, Figure 3). Two additional SNPs, rs17446324 (OR = 0.41; p = 5.00 x 10^−6^) located in the Class 3 Semaphorin (*SEMA3E*) gene on chromosome 7, and rs67731056 (OR = 1.88; p = 6.70 x 10^−6^) in the Insulin Receptor (*INSR*) gene region on chromosome 19, were suggestively associated with asthma risk. Of note, of the four identified associations, the minor allele for only rs17446324 was protective; in the other three loci, the minor allele conferred increased risk for asthma.

**Table 2.** Significant and Suggestive Allelic Associations with Asthma in SAGE II.

| Chromosome | Gene | SNP | A1<sup>1</sup> | A2<sup>2</sup> | MAF<sup>3</sup> | Odds Ratio<sup>4</sup><br>(95% Confidence Interval) | P-Value |
| --- | --- | --- | --- | --- | --- | --- | --- |
| 10 | *PTCHD3* | **rs660498** | **T** | **C** | **0.46** | **1.62 (1.35 – 1.95)** | **2.20 x 10<sup>-7</sup>** |
|  |  | rs2484180 | A | C | 0.49 | 1.54 (1.28 – 1.84) | 2.84 x 10<sup>-6</sup> |
| 7 | *SEMA3E* | rs17446324 | G | A | 0.05 | 0.41 (0.28 – 0.60) | 5.00 x 10<sup>-6</sup> |
| 19 | *INSR* | rs67731056 | C | T | 0.14 | 1.88 (1.43 – 2.48) | 6.70 x 10<sup>-6</sup> |

SNPs in **BOLD** remained significant after correction for multiple testing; Population Specific Genome-wide Threshold: $4.3 \times 10^{-7}$ .
<sup>1</sup>A1: Allele 1 (minor allele)
<sup>2</sup>A2: Allele 2 (major allele)
<sup>3</sup>MAF: Minor Allele Frequency
<sup>4</sup> Adjusted for age, sex, global African Ancestry. P-values reflect the additive effect of the minor allele.

These four SNPs were then evaluated using mediation analysis to determine if the association signal detected was a result of a direct effect of SNP genotype on asthma status, or reflective of an indirect effect on asthma status mediated through obesity. Mediation analysis did not uncover any significant mediation effects via obesity status and provided strong evidence that SNP effects were truly indicative of “direct” asthma associations (Online Resource 2 (Table S1)).

### Replication of Previously Identified Asthma Variants

To determine what proportion of asthma genetic associations discovered in other populations were generalizable to a pediatric African American population, we attempted to replicate previous findings using our study population (Table 3, Online Resource 2 (Table S4)). Of the 59 SNPs evaluated, only three nominally replicated in our study (p < 0.05) (Table 3). Importantly, only a single SNP, rs204993, on chromosome six displayed the same direction of effect in published studies as in our study population.

**Table 3.** Replication of Previously Identified GWAS Associations with Asthma/Asthma-related Phenotypes in SAGE II.

| Chromosome | Gene | SNP | Original OR | Original P-value | SAGE II OR | SAGE II P-Value |
| --- | --- | --- | --- | --- | --- | --- |
| 6 | *PBX2* | rs204993 | 1.17 | 2.00E-15 | 1.23<sup>1</sup> | 0.03 |
| 8 | *SLC30A8* | rs3019885<sup>†</sup> | 1.34 | 5.00E-13 | 0.79<sup>2</sup> | 0.05 |
| 8 | *intergenic* | rs7006742<sup>†</sup> | 0.78 | 1.00E-07 | 0.70<sup>3</sup> | 0.04 |

Only successfully replicated associations (nominal replication threshold ( $p \leq 0.05$ )) are presented above.
<sup>1</sup> Replication was in the same direction as the original study.
<sup>2</sup> Replication was in the opposite direction as the original study.
<sup>3</sup> The original study did not report the reference allele so concordance of effect direction could not be determined.
<sup>†</sup>SNP was imputed in study population
Abbreviations: OR: Odds Ratio

## Discussion

We performed a genome-wide association analysis of asthma susceptibility in African American children and identified novel, and possibly ethnic-specific, genetic associations. Our two most significant associations were between increased asthma susceptibility and genetic variation in the *PTCHD3* gene region; specifically, rs660498, a variant located within 40kb of the *PTCHD3* gene boundary, and rs2484180, a missense SNP responsible for a cysteine to glycine change, located in the first exon of *PTCHD3.* Located less than 50 kb apart, these two variants are in moderate linkage disequilibrium (LD) (r^2^=0.44, p < 0.001). Notably, rs2484180 is located proximally to an exon splice region and is predicted to affect protein stability (Xu and Taylor 2009), and acts as an expression quantitative trait locus (eQTL) in adipose tissues (Ward and Kellis 2012). SNP rs660498 is located in an enhancer histone mark in monocytes and CD14+ cells, acting as an eQTL in whole blood (Ward and Kellis 2012). A recent study identified SNPs within Patched-Chain Domain Protein 1, *PTCHD1,* an important paralog of *PTCHD3,* as predictive of asthma exacerbations (Xu et al. 2011). *PTCHD3* and *PTCHD1* are both predicted to play a role in the Hedgehog signaling pathway, although not as well characterized as *PTCHD1* (Furmanski et al. 2013; Ghahramani Seno et al. 2011). Functional studies highlight the major role that *PTCHD1* plays in modulating T-cell differentiation via regulation of the hedgehog signaling pathway (Furmanski et al. 2013). *PTCHD1* up-regulation was shown to induce a Th2 phenotype in peripheral CC4+ T-cells, and several studies have reported the importance of Th2 cells in the pathophysiology of asthma (Barnes 2001; Fahy 2015; Lloyd and Hessel 2010). To our knowledge, our report is the first instance of genetic variation in the *PTCHD3* gene that is associated with increased asthma susceptibility. Interestingly, this gene has been previously associated with BMI in The Family Heart Study (FamHS) (National Center for Biotechnology Information 2009) and with an interaction of fasting glucose-related traits with BMI (Manning et al. 2012).

We also discovered two suggestively associated non-synonymous SNPs located in the *SEMA3E* and *INSR* genes. Genetic variation in *SEMA3E* has been previously associated with adipose tissue inflammation; it has been hypothesized that inflammatory cells in adipose tissue may promote an inflammatory state in the lungs and encourage the development of asthma (Mohanan et al. 2014; Shimizu et al. 2013). Lastly, *INSR* is the main regulator of insulin, which is known to affect smooth muscle contraction in the lungs and therefore may play a plausible biological role in asthma susceptibility (Schaafsma et al. 2007; Singh et al. 2013). Similar to both *PTCHD3* and *SEMA3E, INSR* has also been linked with several BMI-related phenotypes (Pankov Iu 2013; Parekh et al. 2015).

The discovery that all of our associated variants were located within gene regions previously associated with obesity or BMI-related phenotypes motivated us to investigate whether our identified associations between SNP genotype and asthma status were mediated by obesity status. We performed mediation analysis to determine whether our significant associations were due to a direct relationship between SNP genotype and asthma susceptibility or were in fact the result of an indirect association with asthma susceptibility mediated by obesity/BMI. Results from our mediation analysis provided strong evidence that the effects of all of four of our SNPs of interest, either significantly or suggestively associated with asthma, were independent of obesity status (Online Resource 2 (Table S1)).

We have previously demonstrated that genetic and pharmacogenetic associations for asthma vary among different racial and ethnic groups (Drake et al. 2014; Galanter et al. 2014; Pino-Yanes et al. 2015; Torgerson et al. 2011). Although there are many potential explanations for these observations, one possibility is that there are unique racial and ethnic-specific genetic risk factors for asthma. In this study, we were unable to replicate the majority of prior findings of genetic associations with asthma (Online Resource 2 (Table S2, Table S4)). There are several possible reasons for our inability to replicate previous findings. One possible explanation is that previously discovered associations were population-specific and not generalizable to an African American population. Another possible explanation may be due in part to the differing LD patterns displayed by genetic variants between racial/ethnic groups (Shifman et al. 2003; Xu et al. 2007). It is also possible that a portion of the previous associations identified in adult studies of asthma susceptibility were age-dependent and may not have the same impact in a pediatric cohort (Castro-Giner et al. 2010; Pino-Yanes et al. 2013). We must also consider the possibility that the original findings reported in published GWAs studies were false positives or spurious associations. Finally, we acknowledge that while SAGE II is the largest and most comprehensive pediatric gene-environment study of African American children with and without asthma to date, we may be underpowered to detect small effect sizes due to the sample size.

The identification of novel ethnic-specific genetic variation directly associated with asthma in genes that have been previously associated with obesity and/or adiposity are consistent with the observation of a phenotypic link between these two complex common disorders. The significant racial/ethnic disparity in prevalence and severity of asthma and obesity suggests not only the presence of ethnic-specific factors influencing both diseases, but indicates that there may be ethnicity-specific pleiotropic effects as well. Such findings underscore the importance of studying multiple racial and ethnic groups in genetic association studies. To our knowledge, studies of pleiotropy between asthma and obesity genetics have only been performed in pediatric populations of European descent (Gonzalez et al. 2014; Melen et al. 2013; Melen et al. 2010; Wang et al. 2015). This presents a gap in our scientific knowledge that limits the standard of care available for populations, such as African Americans, who are among the most severely affected by the co-occurrence of these conditions. Our findings are in tune with President Obama’s announcement of the Precision Medicine Initiative, which will include enrollment of at least one million Americans (National Institutes of Health 2015). The goal of this Initiative is to begin a new era of biomedical research that centers on the inclusion of study populations for common disease reflecting the demographic and genetic diversity of U.S. patient populations. As citizens and scientists, we must leverage our skills to ensure that our research furthers this goal and moves us closer to the development of therapies that will improve care in all patients.

## Acknowledgments

The authors acknowledge the patients, families, recruiters, health care providers and community clinics for their participation. In particular, the authors thank Sandra Salazar for her support as the SAGE II studycoordinator. The authors also acknowledge the Lowell Science Research Program, Lowell High School, San Francisco, CA. This work was supported in part by the Sandler Foundation, the American Asthma Foundation, the RWJF Amos Medical Faculty Development Program and the National Institutes of Health (ES015794 and MD006902).

## References

Acock AC (2014) A Gentle Introduction to Stata. Stata Press, College Station, TX.

Akhabir L, Sandford AJ (2011) Genome-wide association studies for discovery of genes involved in asthma. Respirology 16:396–406

Akinbami LJ, Moorman JE, Simon AE, Schoendorf KC (2014) Trends in racial disparities for asthma outcomes among children 0 to 17 years, 2001-2010. J Allergy Clin Immunol 134:547–553 e5

Alexander DH, Novembre J, Lange K (2009) Fast model-based estimation of ancestry in unrelated individuals. Genome Res 19:1655–64

Barnes PJ (2001) Th2 cytokines and asthma: an introduction. Respir Res 2:64–5

Borrell LN, Nguyen EA, Roth LA, Oh SS, Tcheurekdjian H, Sen S, Davis A, Farber HJ, Avila PC, Brigino-Buenaventura E, Lenoir MA, Lurmann F, Meade K, Serebrisky D, Rodriguez-Cintron W, Kumar R, Rodriguez-Santana JR, Thyne SM, Burchard EG (2013) Childhood obesity and asthma control in the GALA II and SAGE II studies. Am J Respir Crit Care Med 187:697–702

Burchard EG (2014) Medical research: Missing patients. Nature 513:301–2

Burchard EG, Oh SS, Foreman MG, Celedon JC (2015) Moving toward true inclusion of racial/ethnic minorities in federally funded studies. A key step for achieving respiratory health equality in the United States. Am J Respir Crit Care Med 191:514–21

Burdett TE, Hall, P.N. (NHGRI), Hasting, E. (EBI), Hindorff, L.A. (NHGRI), Junkins, H.A. (NHGRI), Kiemm, A.K. (NHGRI), MacArthur J. (EBI), Manolio, T.A. (NHGRI), Morales, J. (EBI), Parkinson, H. (EBI), and Welter, D. (EBI) (2014) The NHGRI-EBI Catalog of published genome-wide association studies. www.ebi.ac.uk/gwas. Accessed February 3 2016

Bustamante CD, Burchard EG, De la Vega FM (2011) Genomics for the world. Nature 475:163–5

Castro-Giner F, de Cid R, Gonzalez JR, Jarvis D, Heinrich J, Janson C, Omenaas ER, Matheson MC, Pin I, Anto JM, Wjst M, Estivill X, Kogevinas M (2010) Positionally cloned genes and agespecific effects in asthma and atopy: an international population-based cohort study (ECRHS). Thorax 65:124–31

Center for Disease Control and Prevention (2014) How is BMI Calculated? http://www.cdc.gov/healthyweight/assessing/bmi/adult_bmi/. Accessed October 4 2015

Chang CC, Chow CC, Tellier LC, Vattikuti S, Purcell SM, Lee JJ (2015) Second-generation PLINK: rising to the challenge of larger and richer datasets. Gigascience 4:7

Delaneau O, Marchini J, Zagury JF (2012) A linear complexity phasing method for thousands of genomes. Nat Methods 9:179–81

Drake KA, Torgerson DG, Gignoux CR, Galanter JM, Roth LA, Huntsman S, Eng C, Oh SS, Yee SW, Lin L, Bustamante CD, Moreno-Estrada A, Sandoval K, Davis A, Borrell LN, Farber HJ, Kumar R, Avila PC, Brigino-Buenaventura E, Chapela R, Ford JG, Lenoir MA, Lurmann F, Meade K, Serebrisky D, Thyne S, Rodriguez-Cintron W, Sen S, Rodriguez-Santana JR, Hernandez RD, Giacomini KM, Burchard EG (2014) A genome-wide association study of bronchodilator response in Latinos implicates rare variants. J Allergy Clin Immunol 133:370–8

Fagnani C, Annesi-Maesano I, Brescianini S, D'Ippolito C, Medda E, Nistico L, Patriarca V, Rotondi D, Toccaceli V, Stazi MA (2008) Heritability and shared genetic effects of asthma and hay fever: an Italian study of young twins. Twin Res Hum Genet 11:121–31

Fahy JV (2015) Type 2 inflammation in asthma‐‐present in most, absent in many. Nat Rev Immunol 15:57–65

Furmanski AL, Saldana JI, Ono M, Sahni H, Paschalidis N, D'Acquisto F, Crompton T (2013) Tissue-derived hedgehog proteins modulate Th differentiation and disease. J Immunol 190:2641–9

Galanter JM, Gignoux CR, Torgerson DG, Roth LA, Eng C, Oh SS, Nguyen EA, Drake KA, Huntsman S, Hu D, Sen S, Davis A, Farber HJ, Avila PC, Brigino-Buenaventura E, LeNoir MA, Meade K, Serebrisky D, Borrell LN, Rodriguez-Cintron W, Estrada AM, Mendoza KS, Winkler CA, Klitz W, Romieu I, London SJ, Gilliland F, Martinez F, Bustamante C, Williams LK, Kumar R, Rodriguez-Santana JR, Burchard EG (2014) Genome-wide association study and admixture mapping identify different asthma-associated loci in Latinos: the Genes-environments & Admixture in Latino Americans study. J Allergy Clin Immunol 134:295–305

Genomes Project C, Abecasis GR, Altshuler D, Auton A, Brooks LD, Durbin RM, Gibbs RA, Hurles ME, McVean GA (2010) A map of human genome variation from population-scale sequencing. Nature 467:1061–73

Ghahramani Seno MM, Kwan BY, Lee-Ng KK, Moessner R, Lionel AC, Marshall CR, Scherer SW (2011) Human PTCHD3 nulls: rare copy number and sequence variants suggest a non-essential gene. BMC Med Genet 12:45

Global Asthma Network (2014) The Global Asthma Report 2014. Auckland, New Zealand

Gonzalez JR, Caceres A, Esko T, Cusco I, Puig M, Esnaola M, Reina J, Siroux V, Bouzigon E, Nadif R, Reinmaa E, Milani L, Bustamante M, Jarvis D, Anto JM, Sunyer J, Demenais F, Kogevinas M, Metspalu A, Caceres M, Perez-Jurado LA (2014) A common 16p11.2 inversion underlies the joint susceptibility to asthma and obesity. Am J Hum Genet 94:361–72

Gorina Y (2012) QuickStats: Asthma* Death Rates, by Race and Age Group - United States, 2007-2009. In (MMWR) MaMWR (ed.). Centers for Disease Control and Prevention

Guerrero AD, Mao C, Fuller B, Bridges M, Franke T, Kuo AA (2016) Racial and Ethnic Disparities in Early Childhood Obesity: Growth Trajectories in Body Mass Index. J Racial Ethn Health Disparities 3:129–37

Hoffjan S, Nicolae D, Ober C (2003) Association studies for asthma and atopic diseases: a comprehensive review of the literature. Respir Res 4:14

Hoffmann TJ, Zhan Y, Kvale MN, Hesselson SE, Gollub J, Iribarren C, Lu Y, Mei G, Purdy MM, Quesenberry C, Rowell S, Shapero MH, Smethurst D, Somkin CP, Van den Eeden SK, Walter L, Webster T, Whitmer RA, Finn A, Schaefer C, Kwok PY, Risch N (2011) Design and coverage of high throughput genotyping arrays optimized for individuals of East Asian, African American, and Latino race/ethnicity using imputation and a novel hybrid SNP selection algorithm. Genomics 98:422–30

Howie B, Marchini J, Stephens M (2011) Genotype imputation with thousands of genomes. G3 (Bethesda) 1:457–70

Howie BN, Donnelly P, Marchini J (2009) A flexible and accurate genotype imputation method for the next generation of genome-wide association studies. PLoS Genet 5:e1000529

Imai K, Keele L, Tingley D (2010) A general approach to causal mediation analysis. Psychol Methods 15:309–34

International HapMap C, Altshuler DM, Gibbs RA, Peltonen L, Altshuler DM, Gibbs RA, Peltonen L, Dermitzakis E, Schaffner SF, Yu F, Peltonen L, Dermitzakis E, Bonnen PE, Altshuler DM, Gibbs RA, de Bakker PI, Deloukas P, Gabriel SB, Gwilliam R, Hunt S, Inouye M, Jia X, Palotie A, Parkin M, Whittaker P, Yu F, Chang K, Hawes A, Lewis LR, Ren Y, Wheeler D, Gibbs RA, Muzny DM, Barnes C, Darvishi K, Hurles M, Korn JM, Kristiansson K, Lee C, McCarrol SA, Nemesh J, Dermitzakis E, Keinan A, Montgomery SB, Pollack S, Price AL, Soranzo N, Bonnen PE, Gibbs RA, Gonzaga-Jauregui C, Keinan A, Price AL, Yu F, Anttila V, Brodeur W, Daly MJ, Leslie S, McVean G, Moutsianas L, Nguyen H, Schaffner SF, Zhang Q, Ghori MJ, McGinnis R, McLaren W, Pollack S, Price AL, Schaffner SF, Takeuchi F, Grossman SR, Shlyakhter I, Hostetter EB, Sabeti PC, Adebamowo CA, Foster MW, Gordon DR, Licinio J, Manca MC, Marshall PA, Matsuda I, Ngare D, Wang VO, Reddy D, Rotimi CN, Royal CD, Sharp RR, Zeng C, Brooks LD, McEwen JE (2010) Integrating common and rare genetic variation in diverse human populations. Nature 467:52–8

Joseph M, Elliott M, Zelicoff A, Qian Z, Trevathan E, Chang JJ (2016) Racial disparity in the association between body mass index and self-reported asthma in children: a population-based study. J Asthma/1–7

Lloyd CM, Hessel EM (2010) Functions of T cells in asthma: more than just T(H)2 cells. Nat Rev Immunol 10:838–48

Manning AK, Hivert MF, Scott RA, Grimsby JL, Bouatia-Naji N, Chen H, Rybin D, Liu CT, Bielak LF, Prokopenko I, Amin N, Barnes D, Cadby G, Hottenga JJ, Ingelsson E, Jackson AU, Johnson T, Kanoni S, Ladenvall C, Lagou V, Lahti J, Lecoeur C, Liu Y, Martinez-Larrad MT, Montasser ME, Navarro P, Perry JR, Rasmussen-Torvik LJ, Salo P, Sattar N, Shungin D, Strawbridge RJ, Tanaka T, van Duijn CM, An P, de Andrade M, Andrews JS, Aspelund T, Atalay M, Aulchenko Y, Balkau B, Bandinelli S, Beckmann JS, Beilby JP, Bellis C, Bergman RN, Blangero J, Boban M, Boehnke M, Boerwinkle E, Bonnycastle LL, Boomsma DI, Borecki IB, Bottcher Y, Bouchard C, Brunner E, Budimir D, Campbell H, Carlson O, Chines PS, Clarke R, Collins FS, Corbaton-Anchuelo A, Couper D, de Faire U, Dedoussis GV, Deloukas P, Dimitriou M, Egan JM, Eiriksdottir G, Erdos MR, Eriksson JG, Eury E, Ferrucci L, Ford I, Forouhi NG, Fox CS, Franzosi MG, Franks PW, Frayling TM, Froguel P, Galan P, de Geus E, Gigante B, Glazer NL, Goel A, Groop L, Gudnason V, Hallmans G, Hamsten A, Hansson O, Harris TB, Hayward C, Heath S, Hercberg S, Hicks AA, Hingorani A, Hofman A, Hui J, Hung J, et al. (2012) A genome-wide approach accounting for body mass index identifies genetic variants influencing fasting glycemic traits and insulin resistance. Nat Genet 44:659–69

McGarry ME, Castellanos E, Thakur N, Oh SS, Eng C, Davis A, Meade K, LeNoir MA, Avila PC, Farber HJ, Serebrisky D, Brigino-Buenaventura E, Rodriguez-Cintron W, Kumar R, Bibbins-Domingo K, Thyne SM, Sen S, Rodriguez-Santana JR, Borrell LN, Burchard EG (2015) Obesity and bronchodilator response in black and Hispanic children and adolescents with asthma. Chest 147:1591–8

McGeachie MJ, Stahl EA, Himes BE, Pendergrass SA, Lima JJ, Irvin CG, Peters SP, Ritchie MD, Plenge RM, Tantisira KG (2013) Polygenic heritability estimates in pharmacogenetics: focus on asthma and related phenotypes. Pharmacogenet Genomics 23:324–8

Melen E, Granell R, Kogevinas M, Strachan D, Gonzalez JR, Wjst M, Jarvis D, Ege M, Braun-Fahrlander C, Genuneit J, Horak E, Bouzigon E, Demenais F, Kauffmann F, Siroux V, Michel S, von Berg A, Heinzmann A, Kabesch M, Probst-Hensch NM, Curjuric I, Imboden M, Rochat T, Henderson J, Sterne JA, McArdle WL, Hui J, James AL, William Musk A, Palmer LJ, Becker A, Kozyrskyj AL, Chan-Young M, Park JE, Leung A, Daley D, Freidin MB, Deev IA, Ogorodova LM, Puzyrev VP, Celedon JC, Brehm JM, Cloutier MM, Canino G, Acosta-Perez E, Soto-Quiros M, Avila L, Bergstrom A, Magnusson J, Soderhall C, Kull I, Scholtens S, Marike Boezen H, Koppelman GH, Wijga AH, Marenholz I, Esparza-Gordillo J, Lau S, Lee YA, Standl M, Tiesler CM, Flexeder C, Heinrich J, Myers RA, Ober C, Nicolae DL, Farrall M, Kumar A, Moffatt MF, Cookson WO, Lasky-Su J (2013) Genome-wide association study of body mass index in 23 000 individuals with and without asthma. Clin Exp Allergy 43:463–74

Melen E, Himes BE, Brehm JM, Boutaoui N, Klanderman BJ, Sylvia JS, Lasky-Su J (2010) Analyses of shared genetic factors between asthma and obesity in children. J Allergy Clin Immunol 126:631-7 e1-8

Mohanan S, Tapp H, McWilliams A, Dulin M (2014) Obesity and asthma: pathophysiology and implications for diagnosis and management in primary care. Exp Biol Med (Maywood) 239:153140

National Center for Biotechnology Information (2009) Database of Genotypes and Phenotypes (dbGAP).

National Library of Medicine, Bethesda, MD National Institutes of Health (2015) Precision Medicine Initiative Cohort Program.

Nieminen MM, Kaprio J, Koskenvuo M (1991) Apopulation-based study of bronchial asthma in adult twin pairs. Chest 100:70–5

Nishimura KK, Galanter JM, Roth LA, Oh SS, Thakur N, Nguyen EA, Thyne S, Farber HJ, Serebrisky D, Kumar R, Brigino-Buenaventura E, Davis A, LeNoir MA, Meade K, Rodriguez-Cintron W, Avila PC, Borrell LN, Bibbins-Domingo K, Rodriguez-Santana JR, Sen S, Lurmann F, Balmes JR, Burchard EG (2013) Early-life air pollution and asthma risk in minority children. The GALA II and SAGE II studies. Am J Respir Crit Care Med 188:309–18

Ober C, Yao TC (2011) The genetics of asthma and allergic disease: a 21st century perspective. Immunol Rev 242:10–30

Oh SS, White MJ, Gignoux CR, Burchard EG (2016) Making Precision Medicine Socially Precise. Take a Deep Breath. Am J Respir Crit Care Med 193:348–50

Pankov Iu A (2013) [Major gene mutations associated with obesity and diabetes mellitus]. Mol Biol (Mosk) 47:38–49

Parekh N, Guffanti G, Lin Y, Ochs-Balcom HM, Makarem N, Hayes R (2015) Insulin receptor variants and obesity-related cancers in the Framingham Heart Study. Cancer Causes Control 26:1189–95

Pino-Yanes M, Corrales A, Cumplido J, Poza P, Sanchez-Machin I, Sanchez-Palacios A, Figueroa J, Acosta-Fernandez O, Buset N, Garcia-Robaina JC, Hernandez M, Villar J, Carrillo T, Flores C (2013) Assessing the validity of asthma associations for eight candidate genes and age at diagnosis effects. PLoS One 8:e73157

Pino-Yanes M, Thakur N, Gignoux CR, Galanter JM, Roth LA, Eng C, Nishimura KK, Oh SS, Vora H, Huntsman S, Nguyen EA, Hu D, Drake KA, Conti DV, Moreno-Estrada A, Sandoval K, Winkler CA, Borrell LN, Lurmann F, Islam TS, Davis A, Farber HJ, Meade K, Avila PC, Serebrisky D, Bibbins-Domingo K, Lenoir MA, Ford JG, Brigino-Buenaventura E, Rodriguez-Cintron W, Thyne SM, Sen S, Rodriguez-Santana JR, Bustamante CD, Williams LK, Gilliland FD, Gauderman WJ, Kumar R, Torgerson DG, Burchard EG (2015) Genetic ancestry influences asthma susceptibility and lung function among Latinos. J Allergy Clin Immunol 135:228–35

Purcell SM, Chang, C.C. (2015) PLINK [Version 1.9].

R Development Core Team (2010) R: A language and environment for statistical computing. R Foundation for Statistical Computing, Vienna, Austria

Schaafsma D, Gosens R, Ris JM, Zaagsma J, Meurs H, Nelemans SA (2007) Insulin induces airway smooth muscle contraction. Br J Pharmacol 150:136–42

Sheehan WJ, Phipatanakul W (2015) Difficult-to-control asthma: epidemiology and its link with environmental factors. Curr Opin Allergy Clin Immunol 15:397–401

Shifman S, Kuypers J, Kokoris M, Yakir B, Darvasi A (2003) Linkage disequilibrium patterns of the human genome across populations. Hum Mol Genet 12:771–6

Shimizu I, Yoshida Y, Moriya J, Nojima A, Uemura A, Kobayashi Y, Minamino T (2013) Semaphorin3Einduced inflammation contributes to insulin resistance in dietary obesity. Cell Metab 18:491–504

Singh S, Prakash YS, Linneberg A, Agrawal A (2013) Insulin and the lung: connecting asthma and metabolic syndrome. J Allergy (Cairo) 2013:627384

Sobota RS, Shriner D, Kodaman N, Goodloe R, Zheng W, Gao YT, Edwards TL, Amos CI, Williams SM (2015) Addressing population-specific multiple testing burdens in genetic association studies. Ann Hum Genet 79:136–47

StataCorp. (2011) Stata Statistical Software: Release 12. StataCorp LP., College Station, TX.

Thakur N, Oh SS, Nguyen EA, Martin M, Roth LA, Galanter J, Gignoux CR, Eng C, Davis A, Meade K, LeNoir MA, Avila PC, Farber HJ, Serebrisky D, Brigino-Buenaventura E, Rodriguez-Cintron W, Kumar R, Williams LK, Bibbins-Domingo K, Thyne S, Sen S, Rodriguez-Santana JR, Borrell LN, Burchard EG (2013) Socioeconomic status and childhood asthma in urban minority youths. The GALA II and SAGE II studies. Am J Respir Crit Care Med 188:1202–9

Thomsen SF, Ulrik CS, Kyvik KO, Sorensen TI, Posthuma D, Skadhauge LR, Steffensen I, Backer V (2007) Association between obesity and asthma in a twin cohort. Allergy 62:1199–204

Tingley D, Yamamato, T., Hirose, K., Keele, L., Imai, K. (2014) mediation: R Package for Causal Mediation Analysis. Journal of Statistical Software 59:1–38

Torgerson DG, Ampleford EJ, Chiu GY, Gauderman WJ, Gignoux CR, Graves PE, Himes BE, Levin AM, Mathias RA, Hancock DB, Baurley JW, Eng C, Stern DA, Celedon JC, Rafaels N, Capurso D, Conti DV, Roth LA, Soto-Quiros M, Togias A, Li X, Myers RA, Romieu I, Van Den Berg DJ, Hu D, Hansel NN, Hernandez RD, Israel E, Salam MT, Galanter J, Avila PC, Avila L, Rodriquez-Santana JR, Chapela R, Rodriguez-Cintron W, Diette GB, Adkinson NF, Abel RA, Ross KD, Shi M, Faruque MU, Dunston GM, Watson HR, Mantese VJ, Ezurum SC, Liang L, Ruczinski I, Ford JG, Huntsman S, Chung KF, Vora H, Li X, Calhoun WJ, Castro M, Sienra-Monge JJ, del Rio-Navarro B, Deichmann KA, Heinzmann A, Wenzel SE, Busse WW, Gern JE, Lemanske RF, Jr., Beaty TH, Bleecker ER, Raby BA, Meyers DA, London SJ, Mexico City Childhood Asthma S, Gilliland FD, Children's Health S, study H, Burchard EG, Genetics of Asthma in Latino Americans Study SoG-E, Admixture in Latino A, Study of African Americans AG, Environments, Martinez FD, Childhood Asthma R, Education N, Weiss ST, Childhood Asthma Management P, Williams LK, Study of Asthma P, Pharmacogenomic Interactions by R-E, Barnes KC, Genetic Research on Asthma in African Diaspora S, Ober C, Nicolae DL (2011) Meta-analysis of genomewide association studies of asthma in ethnically diverse North American populations. Nat Genet 43:887–92

Turner SD (2014) qqman: an R package for visualizing GWAS results using Q-Q and manhattan plots. biorXiv

Wang D (2000) sg154: Confidence Intervals for the ratio of two binomial proportions by Koopman's method. Stata Technical Bulletin, 58 edn. Stata Press, College Station, TX.

Wang L, Murk W, DeWan AT (2015) Genome-Wide Gene by Environment Interaction Analysis Identifies Common SNPs at 17q21.2 that Are Associated with Increased Body Mass Index Only among Asthmatics. PLoS One 10:e0144114

Ward LD, Kellis M (2012) HaploReg: a resource for exploring chromatin states, conservation, and regulatory motif alterations within sets of genetically linked variants. Nucleic Acids Res 40:D930–4

Welter D, MacArthur J, Morales J, Burdett T, Hall P, Junkins H, Klemm A, Flicek P, Manolio T, Hindorff L, Parkinson H (2014) The NHGRI GWAS Catalog, a curated resource of SNP-trait associations. Nucleic Acids Res 42:D1001–6

Xu M, Tantisira KG, Wu A, Litonjua AA, Chu JH, Himes BE, Damask A, Weiss ST (2011) Genome Wide Association Study to predict severe asthma exacerbations in children using random forests classifiers. BMC Med Genet 12:90

Xu S, Huang W, Wang H, He Y, Wang Y, Wang Y, Qian J, Xiong M, Jin L (2007) Dissecting linkage disequilibrium in African-American genomes: roles of markers and individuals. Mol Biol Evol 24:2049–58

Xu Z, Taylor JA (2009) SNPinfo: integrating GWAS and candidate gene information into functional SNP selection for genetic association studies. Nucleic Acids Res 37:W600–5

